# Procrustean pseudo-landmark methods in Python to measure massive quantities of leaf shape data

**DOI:** 10.1101/2025.08.08.669192

**Authors:** Asia Hightower, Svea Hall, Ricardo Urquidi Camacho, Alexandra Papamichail, Evan Adamski, Claudia Colligan, Aidan Deneen, Gabriel Dunn, Justine Haziza, Claire Henley, Arnan Pawawongsak, Lachlann Simms, Sean Ward, Manica Balant, Christopher Blackwood, Charles Cannon, Andrea Case, Aman Husbands, Emily B Josephs, Zoë Migicovsky, Rachel Naegele, Eric Patterson, Yenny Alejandra Saavedra-Rojas, Daniel H Chitwood

## Abstract

**Premise:** When examining leaf shapes that are different from one another, it can be difficult to compare both the overall leaf shape and points along the leaf margin in biologically and statistically meaningful ways.

**Method:** To address this problem, we present a simple and user-friendly leaf shape analysis in Jupyter Notebook and Python that uses pseudo-landmarks and Generalized Procrustes Analysis to measure and compare the shape of any leaf. To demonstrate our analysis, we created a repository of real leaves gathered from eight experimental datasets.

**Results:** Using our leaf repository, we explain how we can use pseudo-landmarks to compare all leaf shapes both within and between species using dimension reduction techniques like Principal Component Analysis and can predict leaf shapes using pseudo-landmarks through Linear Discriminant Analysis. Our leaf shape analysis also maps differences in shape as leaves grew around a rosette, showing the transition of shape across development (phyllotaxy). Finally, we showed how we can investigate the relationship between leaf shape variation and genetic diversity by combining shape with genetic data.

**Discussion:** Through the use of Generalized Procrustes Analysis and pseudo-landmarks, our leaf shape analysis presents a powerful tool for examining the shape of any leaf across multiple biological, ecological, evolutionary, and developmental scales.

## Introduction

One of the most basic yet important aspects of research in biology is the phenomenon of shape variation (Thompson, 1963). In the plant sciences, studies commonly measure differences in flower, root, shoot, and leaf shapes within species, across time, space, and other biologically relevant axes (Niklas, 1994). Traditional analyses of shape variation include measuring traits that directly describe shape and size as features, measuring allometric relationships between length, width, and area. Despite limited sampling, manual measurements of leaf shapes provide foundational insights into developmental genetics (Hammond, 1941), paleobotanical responses to climate (Bailey and Sinnott, 1915), and the ability to discriminate between closely related groups for taxonomic purposes (Galet, 1979), demonstrating the power of quantitatively analyzing shape variation. Before the advent of the digital and computational world, these traits were measured by hand. Depending on the shape and size, these measurements can be time consuming and complex, potentially limiting the number of measurable samples.

In the past few decades, we have entered a world of big data science and unprecedented computational power. Today, the shape and size of many plant organs can be automatically annotated using computer vision and measured using a suite of morphometric approaches (Bucksch et al., 2017; Amézquita et al., 2020). Geometric morphometric techniques include landmark-based approaches (Bookstein, 1991) and elliptical Fourier descriptor (Kuhl and Giardina, 1982) analysis. For landmark-based techniques, closed contours are aligned against each other using corresponding coordinates that minimize a Procrustes distance through the functions of translation, rotation, scaling, and reflection (Gower, 1975). True landmarks are biologically homologous points and often, landmark-based analyses are used to measure and describe important floral landmarks including lateral and modified petals, in addition to measuring symmetry and the angles of sepals, petals, and other reproductive organs (Savriama, 2018). For leaf landmarks, different groups may have a similar number of lobes, leaflets, or terminal points in addition to other leaf margin features. One example of landmarking in leaves is ampelography, the quantitative study of grapevine leaf shape, in which landmark approaches take advantage of the fact that all grapevine leaves have five major veins, creating a homologous coordinate system between leaves (Chitwood et al., 2014). Therefore, when biologically homologous points are present, landmark-based analyses are exceptionally powerful, in that the biological and statistical measures of shapes correspond to each other.

Landmarks may not always be possible between shapes that include non-homologous points or not of the desired resolution to capture many fine leaf margin features, depending on the biological context that informs homology. For lateral organs, traits like the number, angle, size, and location of lobes or leaflets may change between species or across time and space (Balant et al., 2024). Unfortunately, the lack of homology may make comparing leaf shapes difficult-to-impossible both computationally and biologically, or limit the measurement of subtle features, incompletely quantifying forms readily apparent by eye. When homology is otherwise not present or is difficult to describe across the lamina, leaf shapes can be compared using equidistantly placed pseudo-landmarks (Dujardin et al., 2014) between the base and tip points. Using both landmarks and pseudo-landmarks, in a Procrustean analysis (Gower, 1975; Schindelin et al., 2012), shapes are translated, scaled, rotated, and flipped to minimize the overall distance between two shapes. In generalized Procrustes analysis (GPA), we calculate a Procrustes distance and superimpose all shapes to an arbitrary sample, generate a new mean from the superimposed shapes, and repeat until the Procrustes distance between two successive mean shapes is below an arbitrarily low threshold. Once a mean shape has been calculated by GPA, all samples can be superimposed to allow for morphometric analysis. By mapping the dissimilarity between shapes based on landmark/pseudo-landmarks, we can create continuous representations of shape across a large number of shapes. Similar to geometric morphometrics, elliptical Fourier descriptor analysis (EFA) uses landmarks/pseudo-landmarks to describe curves. In EFA, sine and cosine functions are alternatively used in a harmonic series to describe a closed curve or outline.

Geometric morphometric, GPA, and EFA techniques have been included in multiple packages across programs to examine specific kinds of shapes, allowing for the analysis and generation of large shape datasets. In R, popular shape analysis packages include shapes (Dryden and Mardia, 2016) for Procrustes analysis and MOMOCS (Bonhomme et al., 2014) is used to compare and visualize open and closed outlines using images and EFA. In Python, contours are often extracted from images using the program OpenCV (Bradski, 2000). Additionally, many shape analyses can be completed entirely in ImageJ/FIJI (Schindelin et al., 2012), where shapes can be measured, outlined, and extracted for further analysis using a multitude of 2D and 3D shape related plug-ins. Comparatively, the rise in studies investigating leaf shape variation has led to the development of a number of leaf-specific image identification and segmentation programs. Widely used programs like LAMINA (Bylesjö et al., 2008), Morpholeaf (Biot et al., 2016), MowJoe (Failmezger et al., 2018), LeafletAnalyzer (Liao et al., 2017), and MuLES (Bowman et al., 2023) use image recognition, automated segmentation, and classification to compare leaf shape variation. However, often due to the “ready-to-use” nature of these programs, they are limited in their degree of customization, limiting the number and types of non-model plant species that can be analyzed. Therefore, as the context of plant shapes and desired analyses continue to increase across image types, species, and timepoints, the need for new and flexible analysis methods only continues to grow.(Liao et al., 2017; Failmezger et al., 2018)(Bylesjö et al., 2008; Bowman et al., 2023)

Here, we present methods for comparing and describing differences in leaf shapes with or without homology using geometric morphometric methods coded in python, presented in accessible and reproducible Jupyter notebooks (Kluyver et al., 2016). We address these issues by using the building blocks already available for studying leaf shape variation and applying pseudo-landmarks and Procrustes analysis in python. Our leaf shape analysis is based on the fact that all leaves contain comparable homologous points at their tip and base. In this study, we present a fast, simple, and cost-effective method that is customizable and usable with almost any leaf shape. We demonstrate how our pseudo-landmarks and Procrustean leaf shape analysis can quickly measure large numbers of leaves to create theoretical representations of leaf shape, visualize relationships through dimension reduction techniques, and present continuous representations of genetic and developmental effects on leaf shape using genetic and developmental models. We use an ensemble of datasets from a diverse group of study species collected across the globe in combination with previously published data to demonstrate the effectiveness of pseudo-landmarks and Procrustes analysis across a diversity of leaf shapes.

## Methods

### Required and optional data for leaf shape analysis

In this study, we’ve separated data into required and optional information for a given dataset. Required information includes the data necessary for assigning pseudo-landmarks, rotating each leaf, and comparing leaves using pseudo-landmarks during Procrustes analysis. Required data includes a leaf outline - either as a binary image or a text file of the leaf outline as XY Cartesian coordinates. Each leaf also requires the XY coordinates for the tip and base of each leaf. The tip and base landmarks are used to rotate and orient each leaf, with the tip at the top and the base at the bottom. Finally, each leaf requires the number of pixels per centimeter to scale each leaf in the dataset. The pixels per centimeters, noted as “px_cm”, is typically calculated by including a ruler in the leaf image scan and using ImageJ to measure the number of pixels per 1cm. The description for how each dataset calculated px_cm, including the software used, can be found in table one under “leaf imaging and segmentation”. Optional information for each dataset includes information required for identifying demographic, location, and genetic data for each leaf in the dataset. However, the leaf shape analysis can be performed without this information. For each leaf, this information includes the species, population, and genotype data, genetic and transcriptomic data, location data such as latitude, longitude, elevation, temperature, and precipitation, growth conditions, and test treatments (table two). This leaf shape pipeline can be used to examine data along any additional axes to answer a multitude of biological questions. A description of the required data for each leaf can be found in table two and a description of the optional data for each dataset, including plant growth information can be found in table one.

**Table one.** Leaf shape collection and imaging information for all datasets. We collected information on which species were included in each dataset, where geographically leaf material was harvested for each dataset, seed source for species in each dataset, and leaf selection criteria for each dataset. Additionally, we collected the imaging protocol, including equipment used for imaging for each dataset. Finally, we collected the segmentation and landmarking protocol for each dataset, including the final number of leaves.

**Table two.** Individual leaf metdata and measurements. For all leaves in the total dataset, the filename (“file), dataset ID (“dataset”), the length of 1 cm in pixels (“px_cm”), genotype (“genotype”), base coordinates (“base_x” and “base_y”), and tip coordinates (“tip_x” and “tip_y”) were supplied. When applicable, the node (“node”) and rosette ID (“rosette_id”) were also supplied. Additionally, we calculated PC1, PC2, the width, length, area, solidity, asymmetry, circularity (“circ”), and aspect ratio (“ar”) for each leaf using our leaf shape analysis.

### Compiling a unique and diverse set of leaf shapes

We collected eight unique datasets from collaborators from across the globe to test our leaf shape analysis. The final leaf dataset used to test our leaf shape analysis included a total of n = 5129 leaves and n = 48 species. The *Malus* dataset, which includes both wild and domesticated *Malus* species, featured the largest number of leaves (n = 1560) (Migicovsky et al., 2017). After the *Malus* dataset, from highest to lowest in number, our final dataset included *Quercus spp*. (n = 1142), *Arabidopsis thaliana* (n = 771) (AraPheno, n.d., n.d.), *Lobelia spp.* (n = 637), *Cannabis sativa* (n = 340) (Balant et al., 2024), *Beta vulgaris* (n = 269), *Erythroxylum coca* (n = 260), and *Capsella bursa-pastoris* (n = 150) leaves (table one). Three datasets included more than one species: Malus spp. (n = 35), *Lobelia spp.* (n = 10), and *Quercus spp*. (n = 3). Additionally, the leaf images from the *B. vulgaris*, *C. bursa-pastoris*, *A. thaliana*, *Lobelia spp.*, and *C. sativa* datasets were collected from plants grown in a common garden (growth chamber or greenhouse) setting (table one). In contrast, leaf images from the *Malus spp.* dataset were collected from trees grown in two USDA field collection sites (table one). Leaf images from the *Erythroxylum coca* dataset were grown and collected by Colombian national police officers at a garden established by the Anti-Narcotics Directorate (San Luís, Tolima, Colombia) (table one) and leaf images from the *Quercus spp*. dataset were collected from public and privately owned land throughout western Texas and Oklahoma and from herbarium specimens also collected throughout western Texas and Oklahoma from various years (table one). Separate from the above leaves, we analyze 3,299 leaves from *Passiflora/maracuyá spp.* and 11,276 *Solanum spp.* (tomato) leaflets that were previously published (Chitwood et al., 2013; Chitwood and Otoni, 2017a, b) to demonstrate how these methods can be applied to continuous models of leaf development and high-resolution genetic mapping of leaf features, respectively.

### Landmarking a diverse set of leaf shapes

The overall leaf shape analysis includes leaf imaging and segmentation (depending on the dataset), the creation of binary leaf images or XY coordinate leaf outlines, the orientation and scaling of each leaf, comparisons of all leaves using GPA, and further analysis using dimension reduction techniques and developmental or genetic modeling. In this study, each dataset included the required information for the initial analysis in addition to information for further analyses within the dataset. For all datasets, we requested leaf images either as binary images and/or as leaf outlines including XY Cartesian coordinates. Therefore, each research group selected leaf images based on their own criteria for each species (table one). We requested scale information in the form of the number of pixels per centimeter for each leaf image. Each group also provided XY coordinates for the tip and the base of each leaf for orientation. Datasets with species or genotype information also included these as a factor in their metadata. All initial required and optional data can be found in (table two). In our shape analysis, we first analyzed each dataset separately, as described in the pseudo-landmark interpolation section. 100 pseudo-landmarks were placed at equal distances along each leaf margin, creating a leaf outline. Then, we saved an oriented, rescaled, and landmarked array for each dataset and combined all datasets into an array (scaled to centimeters). The combined array was used to perform generalized Procrustes analysis.

### Leaf shape analysis

#### Extracting individual leaves and landmarks

For each dataset, leaves were selected, imaged, and landmarked using ImageJ/FIJI or WinRhizo for *Lobelia spp* leaves. The *B. vulgaris*, *C. sativa*, and *C. bursa-pastoris* datasets all included leaves from the entire plant, including the rosette and cauline leaves when applicable (table one). The Malus spp, *A. thaliana*, *E. coca*., and *Quercus spp.* datasets included a specific number of leaves collected from each plant. The base and then tip of each leaf were manually landmarked in ImageJ using the point selection too. For each leaf, the base and tip were saved as Cartesian (XY) coordinates in a separate file and included in the final metadata comma separated file (csv).

#### Pseudo-landmark interpolation

To compare leaves across multiple species, genotypes, locations, and time, we used pseudo-landmarks, or 100 equidistant points along the leaf margin to compare leaf shapes. In this pipeline, we first uploaded our leaf shape csv and all jpg images into a Jupyter notebook (Kluyver et al., 2016) by dataset. To place pseudo-landmarks around the leaf starting at the tip, all images were read in grayscale and the contour of the largest object, the black leaf, was selected using the package OpenCV (Bradski, 2000). Using the contours function in OpenCV, we extract the XY coordinates and the area of each leaf image. We then use an interpolation function to convert the leaf coordinates to a high-resolution number of points. In this context, the interpolation function is using the known set of real points generated from the leaf contour to estimate 1000 high resolution points for each leaf. We then find the Euclidean distance, or the length of a line segment, between the base and tip landmarks for each leaf and we then index the base and tip points. We then reset the base point to zero and reindex the pseudo-landmarks, creating a single array for all leaf coordinates. We then interpolate the left and right sides of each leaf using the reindexed landmarks to estimate 50 pseudo-landmarks on each side, equaling 100 pseudo-landmarks in total. We then orient each leaf by rotating each leaf upward based on the base and tip landmarks and we rescale each leaf by the pixels per centimeter provided for each leaf image. Finally, we save each oriented and rescaled leaf as an array. For each dataset, we saved a separate array of oriented and rescaled leaves. We created a final array of all oriented and rescaled leaves called the “cm_arr” array. Then we created a final csv called “mdata” which combined all required and optional data for each dataset.

#### Generalized Procrustes analysis (GPA)

To calculate the Procrustes mean, we first select the number of pseudo-landmarks and dimensions. We then calculate the mean shape using the gpa_mean function which includes our array of leaf shapes, the number of pseudo-landmarks, and the number of dimensions. This mean leaf was generated by comparing each leaf to each other leaf by the pseudo-landmarks to create an average of all leaves. We then align all leaves to the newly generated mean leaf to create an array of all aligned leaves. To generate a mean leaf per dataset, we used the same method by substituting the “cm_arr” array with the dataset array.

### Analysis of leaf dimensions

For each leaf, we calculated the width, length, area as centimeters squared, solidity as the ratio of area to convex hull area, circularity as the ratio of area to perimeter, and the aspect ratio - or the ratio of width to length. All values were saved in the mdata csv.

### Principal Component Analysis (PCA)

To perform PCA on all Procrustes aligned leaves using the 100 pseudo-landmarks, we reshaped our Procrustes aligned array from 4D to 2D, calculated our desired number of PCs, executed the PCA function, and saved PC1 and PC2 for each leaf in the original csv. The loadings of this PCA were the combinations of the 100 pseudo-landmarks. We calculated and visualized a morphospace of all leaves in our dataset by using the values of PC1 and PC2 for each leaf to calculate inverse theoretical leaves using the pca.inverse_transform function. We visualized all leaves in PC space by plotting all leaves onto the morphospace as a scatterplot.

### Linear Discriminant Analysis (LDA)

To perform LDA on all leaves in our final dataset, we first created a new dataframe based on the dataset. We imported dataset labels from the mdata csv and included landmark data as X values and the dataset labels as y values. We then fit the LDA model, retrieved the LDA scalings and coefficients, and performed genotype prediction. We then created a confusion matrix based on the true genotype values using the original csv and the predicted genotype values. We then validated our model using Stratified K-fold cross validation scikit-learn package. We used a default K = 5, with 3 repeats and a random state of 42. Finally, we saved the confusion matrix in the new dataset data frame and visualized the confusion matrix comparing true and predicted datasets in addition to all leaves in LD space using LD1 and LD2.

### Modeling leaf shape by relative node – across development

We used a maracuyá (also known as Passiflora) dataset example, including previously published leaf shape data from 40 maracuyá species, to demonstrate the change in leaf shape from plant shoot to plant tip using relative nodes. First we define the number of intervals – or the number of modeled leaves we want to produce. This number depends on the number of nodes included in the dataset. Series of three or more nodes are recommended for modeling. We then find the total number of pseudo-landmarks and the total number of individual plants in the dataset. We then create an array that includes the intervals, number of plants, and number of pseudo-landmarks. After, we create empty lists to store species and morphotype information. Future dataset’s modeled leaves can include any axes information already defined in the dataset. Finally, we model changes in leaf shape across development by first finding the relative node number for each leaf for each individual plant. We then use the polyfit function from the numpy package. This function fits a polynomial to a degree using XY coordinates. The polyfit function includes the relative nodes as x values, the 100 pseudo-landmarks for each node as y values, and the lowest polynomial order (polyorder) possible. For this dataset, we used a polyorder of two. We then modeled the polyfit values over each node, creating 40 modeled leaves with 100 pseudo-landmarks per plant. We then save all modeled leaves as an array and all species and morphotype information as a dataframe. We visualized the 40 modeled leaves per dataset and morphotype by plotting each modeled leaf in a series of increasing node numbers, colored by the morphotype. We then visualize the developmental model for each plant by projecting the intervals for each model back onto the morphospace PCA. We represent each model as a curve, where the individual leaves are connected on the curve and colored by their position in the series. We specify a universal color map for the modeled leaf position in each series using the "viridis" color palette, where yellow represents leaves at or near the base of the plant and purple represents leaves at or near the tip of the plant.

### Modeling leaf shape by relative node – by genetic data

We used a previously published *S. lycopersicum* dataset of leaves from 76 introgression lines (IL), each with small introgressions from *S. pennellii* into a *S. lycopersicum* background. For each introgression, we reindexed each IL to include either a one or zero for the presence or absence of each introgression, tiling across the *S. lycopersicum* genome. Similar to modeling across development, we used pseudo-landmarks to Procrustes align all right later, left lateral, and terminal leaflets in this dataset. After generating a mean leaf for the right, left, and terminal leaves, we calculated the Euclidean distance of each pseudo-landmark to the mean leaflet for each leaflet (noted as displacement). We then calculated the Spearman’s rank correlation coefficient between the displacement for each pseudo-landmark and the presence/absence of an introgression for each IL, generating a p-value. We then visualized the degree of displacement by each pseudo-landmark for its correlated introgression, with each significant correlation colored by its respective p-value.

## Results

### Procrustes analysis reveals relationships between shapes using pseudo-landmarks

Mean leaf comparisons using 100 pseudo-landmarks and generalized Procrustes analysis revealed a consensus leaf that lacks margin features, such as serrations or lobes, due to averaging (Fig. 1A (left)). This averaging out effect occurs when features fail to align with specific pseudo-landmarks between samples. Within each dataset, there is also considerable variation between leaves (Fig. 1B). Comparisons between the overall mean leaf and the mean leaf per dataset shows that each dataset mean leaf is unique (Fig. 1B). The *C. bursa-pastoris*, *Lobelia spp.*, and *Quercus spp*. datasets include leaves that have high, non-homologous intra-dataset variation. Due to this, the differences in shape are averaged out and not visible in the mean leaves for these three datasets. In comparison, the *Malus spp.*, *A. thaliana*, *E. coca,* and *B. vulgaris* leaves resemble real leaves in each dataset and include features like serrations (Fig 1B). In particular, the *C. sativa* mean leaf is a simple palmate leaf, the mean of which has three prominent lobes, resulting from the similar placement of these features across leaves, while the *B. vulgaris* mean leaf is a simple leaf with a broad, sagittate blade and prominent petiole.

**Figure one.**
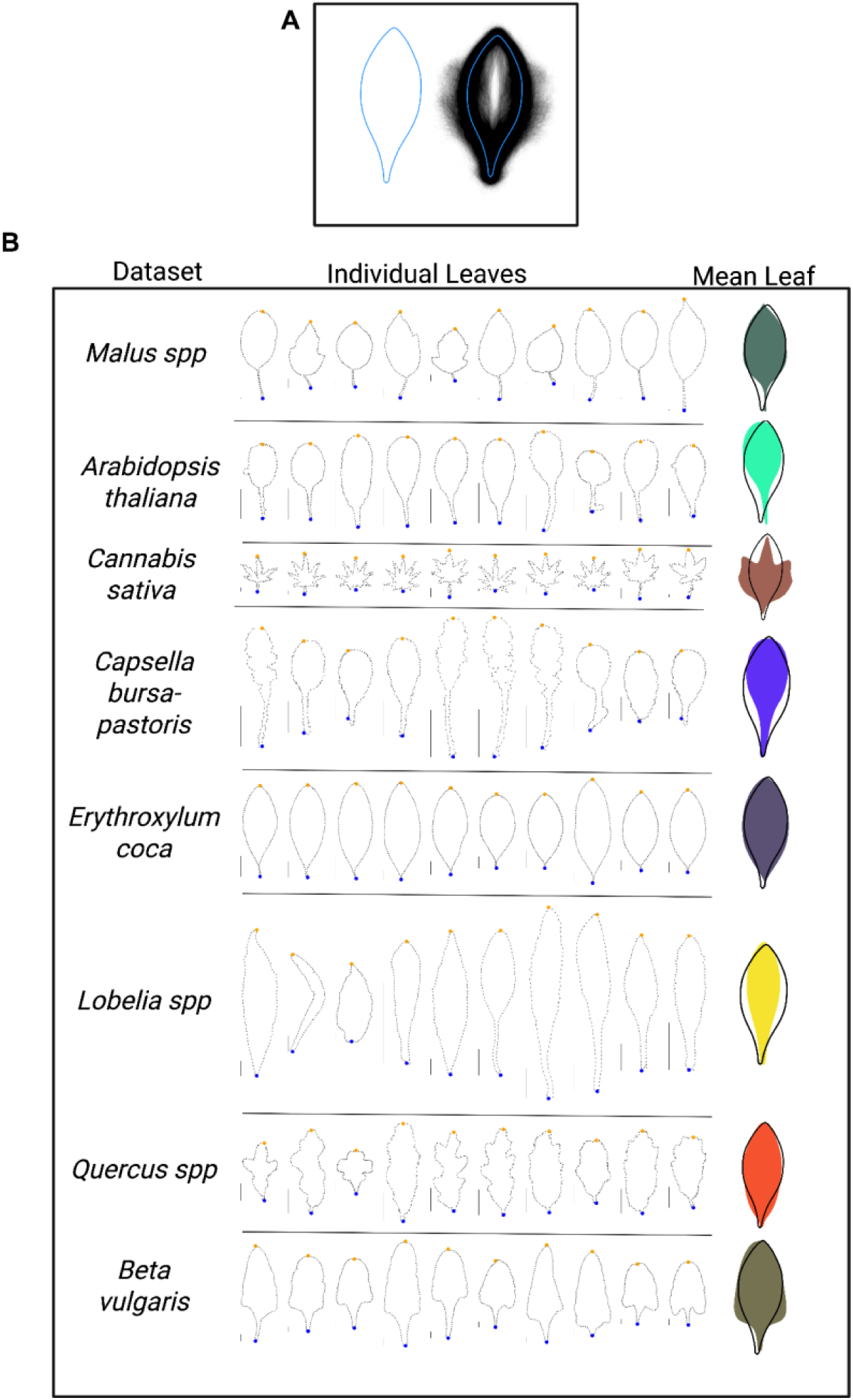
A description of the diversity of leaf shapes across all datasets. **A**. The mean leaf for the total dataset generated through Procrustes alignment (left) and the same mean leaf for the total dataset superimposed onto each individual leaf outline (right). **B**. For each species dataset, an alignment of ten random leaf outlines including 100 pseudo-landmarks extracted from each species dataset are presented. Orange circle = the tip of each leaf. Blue circle = the base of each leaf. Black line = 1 cm scale bar. For each species dataset, the total dataset mean leaf (black) is superimposed onto the colored species mean leaf.

### Shape descriptors reveal relationships between leaves based on total shape

In our analysis, shape can be directly measured using shape descriptors like circularity 4*π*(*area*/*perimeter*^2^). Circularity is a measure of the undulations in shape and is often used to quantify and describe the degree of lobing around a leaf margin. Circularity describes the roundness of leaves in relation to a perfect circle. Therefore, leaves with a lower measure of circularity have a circularity score closer to 0 and are the least round while leaves with a higher measure of circularity are closer to 1 and more round. Circularity values decrease with undulations like lobes and serrations, long tapering petioles, or elongated shapes. The total dataset included an overall circularity range from 0.1 to 0.85 and a mean circularity of 0.45, encompassing the range of all possible shapes (Fig. 2A,2B). When examining the distribution of circularity within each dataset (Fig. 2B), the *C. bursa-pastoris* (Fig. 2C)*, Lobelia spp, Quercus spp, Malus spp,* and *A. thaliana* datasets include the most variation in circularity (between 0.07 to 0.86), followed by *B. vulgaris* and *E. coca* (between 0.41 to 0.85) and finally *C. sativa* (between 0.07 to 0.33) (Fig. ). *C. sativa* leaves exhibit the lowest circularity overall and some of the most lobing. Here, circularity is especially sensitive to the elongated shape and tapering proximal blade present in the *E. coca* leaves (Fig. 2D). Overall, when we consider within-dataset shape variation especially, *C. bursa-pastoris, Lobelia spp, Quercus spp, Malus spp, and A. thaliana* leaves are the least similar to one another while *C. sativa, B. vulgaris, and E. coca leaves* are the most similar to one another.

**Figure two.**
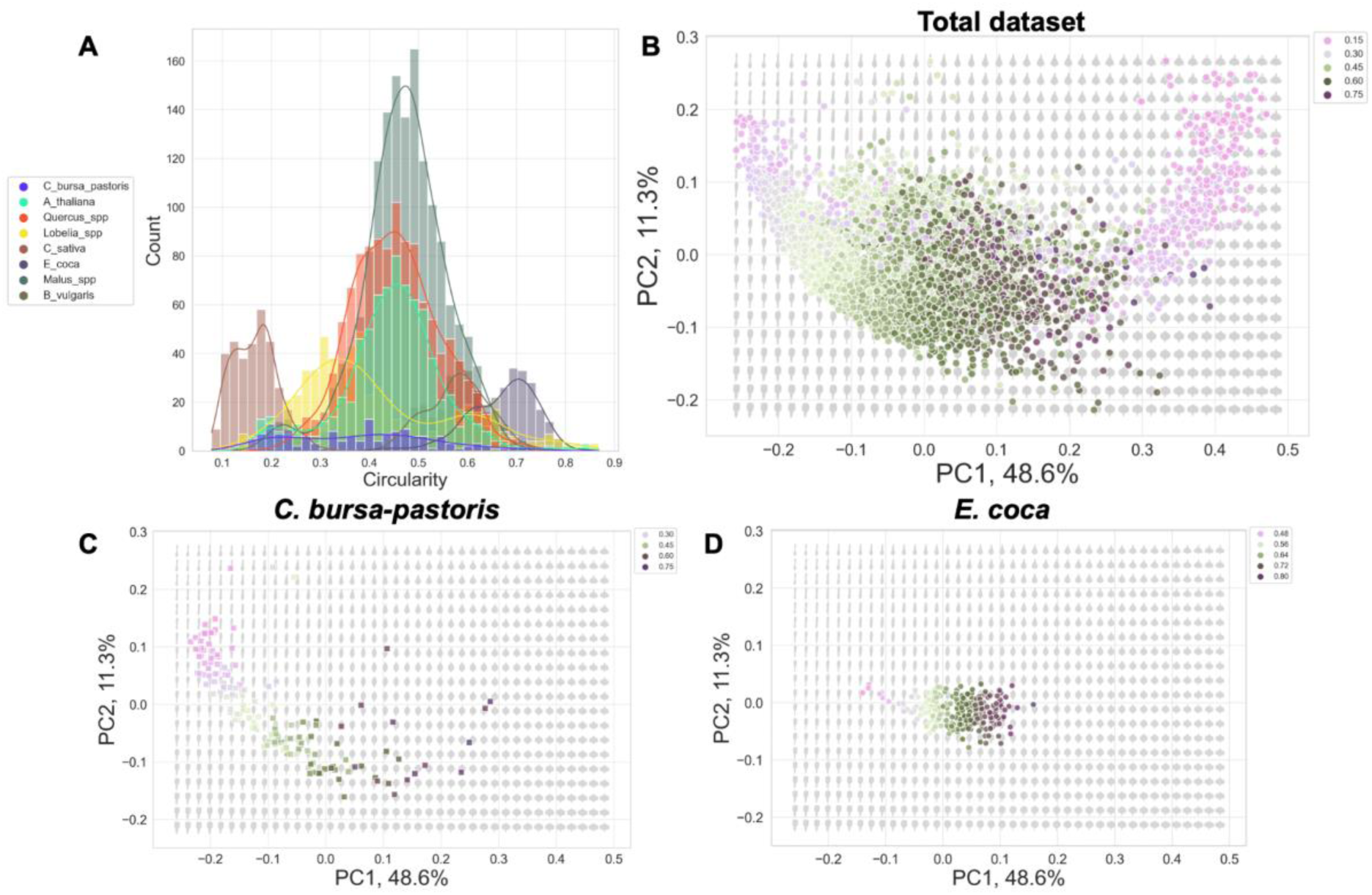
Leaf shape diversity described by circularity. **A**. A histogram of circularity values of leaves in each species dataset. **B**. A morphospace PCA colored by circularity, populated by leaves from the total dataset. **C**. A morphospace PCA colored by circularity, populated by leaves from the *C. bursa-pastoris* dataset. **D**. A morphospace PCA colored by circularity, populated by leaves from the *E. coca* dataset. In each morphospace PCA colored by circularity, leaves are colored along a continuous circularity spectrum, with low circularity as light pink, medium circularity as light green to dark green, and high circularity as dark purple.

### Morphospace PCA reveals relationships between leaves based on parts of the leaf

In comparison to directly measuring total shape using shape descriptors, we can also use pseudo-landmarks with dimension reduction techniques to determine and predict which parts of the leaf margin contribute most to the overall variation between leaves. Through Principal Component Analysis (PCA), we can determine relationships between shapes along multiple axes by examining the overlap in shapes in PC space visualizing a morphospace. In a morphospace PCA, the PC loadings for combinations of pseudo-landmarks can reveal which parts of the leaf margin most contribute to differences between the leaves along each PC axis. Additionally, by performing an inverse PCA, we can recreate the shapes that contribute most to overall differences along PCs. In the morphospace PCA for the total dataset, the width of the leaf changes from left to right along PC1, with thinner leaves on the left and wider leaves, especially those with multiple lobes and leaflets, found on the right edges (Fig. 3). This variation in leaf aspect ratio, captured by a single value in PC1, explains 48.6% of observable variation in our data. Additionally, the amount of proximal lamina and the presence of well-defined petioles changes from top to bottom along PC2. Leaves found at the top of the morphospace do not have well defined petioles and contain the most proximal lamina while leaves at the bottom of the morphospace have well defined petioles and do not contain much proximal lamina (Fig. 3). The distinctness of the petiole captured by PC2 explains 11.3% of the variation in our data.

**Figure three.**
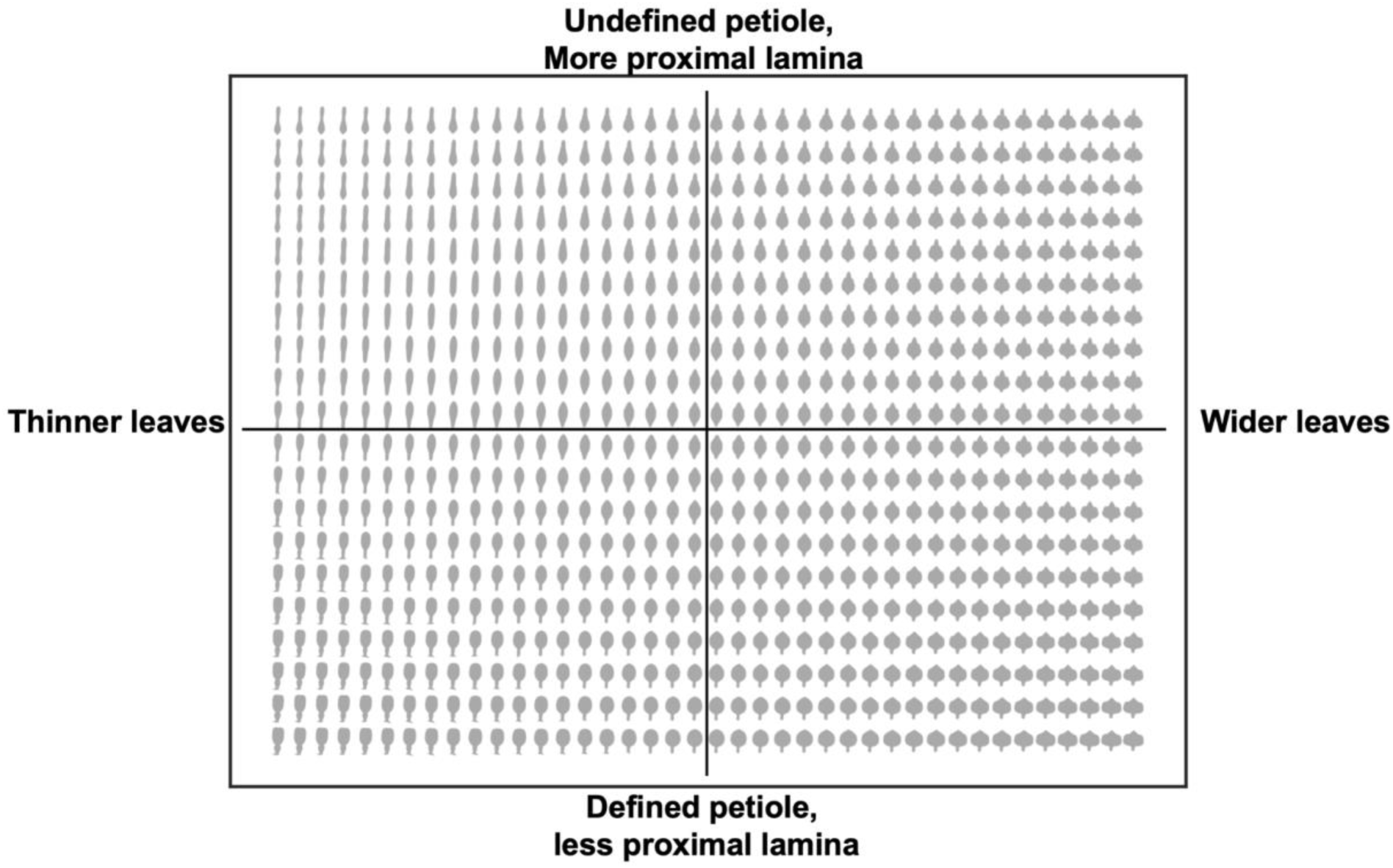
Morphospace of theoretical leaves recreated from the inverse of PC1 and PC2. Four quadrants are defined left to right (width of leaves) and top to bottom (petiole definition, amount of lamina).

The PCA morphospace revealed interesting relationships between our datasets (Fig. 4A). *C. sativa* leaves were separated from all other datasets. Within the morphospace, *C. sativa* occupied the area most represented by leaves with multiple tips, while the other seven datasets occupied the rest of the morphospace, which includes leaves with a large amount of lobing variation. In comparison, the *Quercus spp*. and *Malus spp* leaves formed distinct groups with some overlap (Fig. 4B). Leaves from all remaining datasets showed significant overlap in PC space (Fig. 4B). When we examine the distribution of individual leaves across the morphospace for each dataset, two major trends emerge. Within a dataset, the individual leaves encompass a significant range in PC space, suggesting the variation in leaf shapes within a dataset is large compared to all variation among all datasets. Within their respective datasets, the *Quercus spp*, E. coca, and *B. vulgaris* leaves are more similar to each other than the *Malus spp*, *A. thaliana*, *C. sativa*, *C. bursa-pastoris*, and *Lobelia spp* leaves (Fig. 4B). This would suggest that *Quercus spp*., *E. coca*, and *B. vulgaris* leaves have less variation within their respective datasets along PC1 and PC2. Overall, using a morphospace PCA can be informative for visually delineating relationships between species (or genera, families, etc) based on shape alone.

**Figure four.**
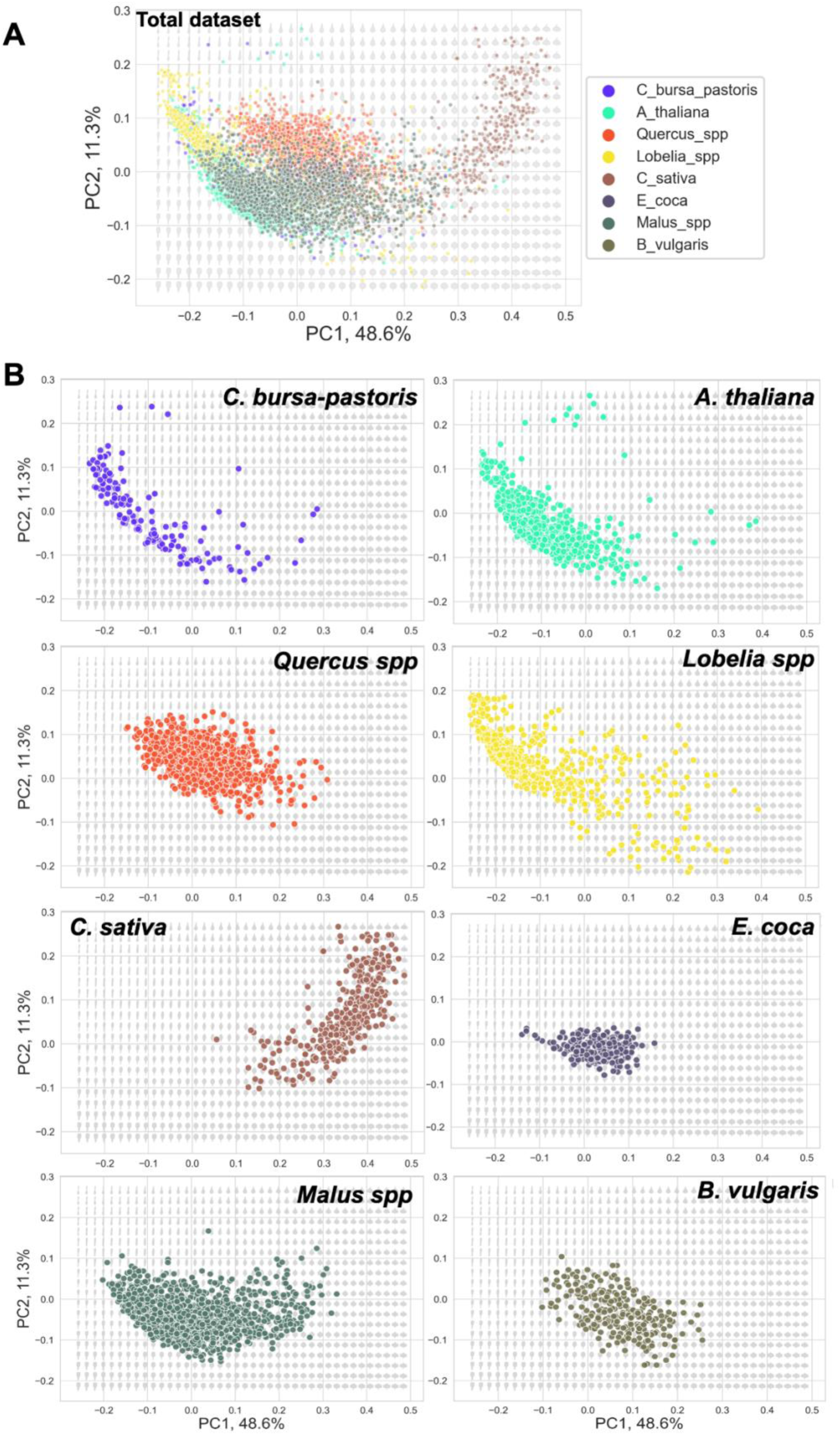
Morphospace PCA of all leaves. The projected morphospace includes eigenleaves created by taking the inverse of the PCA. These eigenleaves were created from the 100 pseudo-landmarks outlining each leaf image. **Top panel:** morphospace PCA of total dataset. **Bottom panels:** Each species dataset morphospace PCA was plotted against the total morphospace. Point colors correspond with each species dataset.

### LDA predicts relationships between leaves along biologically relevant axes

We can use the parts of the leaf most responsible for differences in shape to predict and interpret relationships between leaf shapes along biologically important axes through Linear Discriminant Analysis (LDA). LDA is like PCA, but instead of defining axes based on linear combinations that explain the most variation in the data overall, linear discriminant axes are defined by the linear combination of features most responsible for separating defined groups. Because LD axes are defined by specified groups, it can be used for modeling, in the form of modeling categorical group identity as a function of a number of continuous variables (such as Procrustean-aligned coordinate values).

In using LDA, our goal was to predict which dataset a leaf belonged to using the 100 pseudo-landmarks. Notably, due to the unequal numbers of leaves in each dataset, we used Stratified K-fold cross validation to create a uniform sampling for both our training and testing groups. Stratified sampling helps to improve the likelihood of true positive predictions by ensuring that each dataset is proportionally represented in both the training and testing groups. Without stratified sampling and instead by using random sampling, there is the possibility that datasets with small numbers like the *C. bursa-pastoris* dataset (n = 150) are not represented proportionally.

When we performed the LDA, our dataset achieved a mean accuracy of 87%, with 511 leaves falsely predicted and 4618 leaves out of 5129 leaves correctly predicted. Our confusion matrix shows that the *Malus spp* dataset included the majority of correctly predicted leaves (Fig. 5A). The *C. bursa-pastoris* leaves were the least correctly predicted (n = 67 out of 150) and were often falsely predicted as *A. thaliana* leaves (n = 60) (Fig. 5A). 97% of *E. coca* leaves, 96% of *C. sativa* leaves, 92% of *B. vulgaris* leaves, 91% of *Quercus spp leaves, and* 80% of *Lobelia spp* leaves were correctly predicted in our LDA (Fig. 5A). Similar to the PCs in PCA, the LDs are the linear combinations of our predictor variables, the 100 pseudo-landmarks. When we visualize our prediction results, we see that datasets separate along LD1, with a clear delineation between *C. sativa* leaves and every other dataset (Fig. 5B). Along LD2, while we see distinct separation between *Malus spp* and *Quercus spp* leaves, we also see significant overlap between the remaining datasets in the center of the cluster (Fig. 5B, 5C). While results suggest leaves from most datasets can be reliably predicted using pseudo-landmarks, our model may be sensitive to both the low number of *C. bursa-pastoris* leaves and the similarity between *C. bursa-pastoris* and *A. thaliana* leaf shapes. By combining the results of both dimensional reduction techniques, we are able to model how datasets relate to each other. When we consider the PCA and LDA results together, we see that *C. sativa* leaves consistently separate into a distinct group, *Malus spp* and *Quercus spp* leaves share both PC and LD space but can be delineated into distinct groups, and the remaining datasets consistently show overlap in leaf shape based on both the 100 pseudo-landmarks and circularity. Both PCA and LDA using pseudo-landmarks present important initial information on relationships between and within datasets.

**Figure five.**
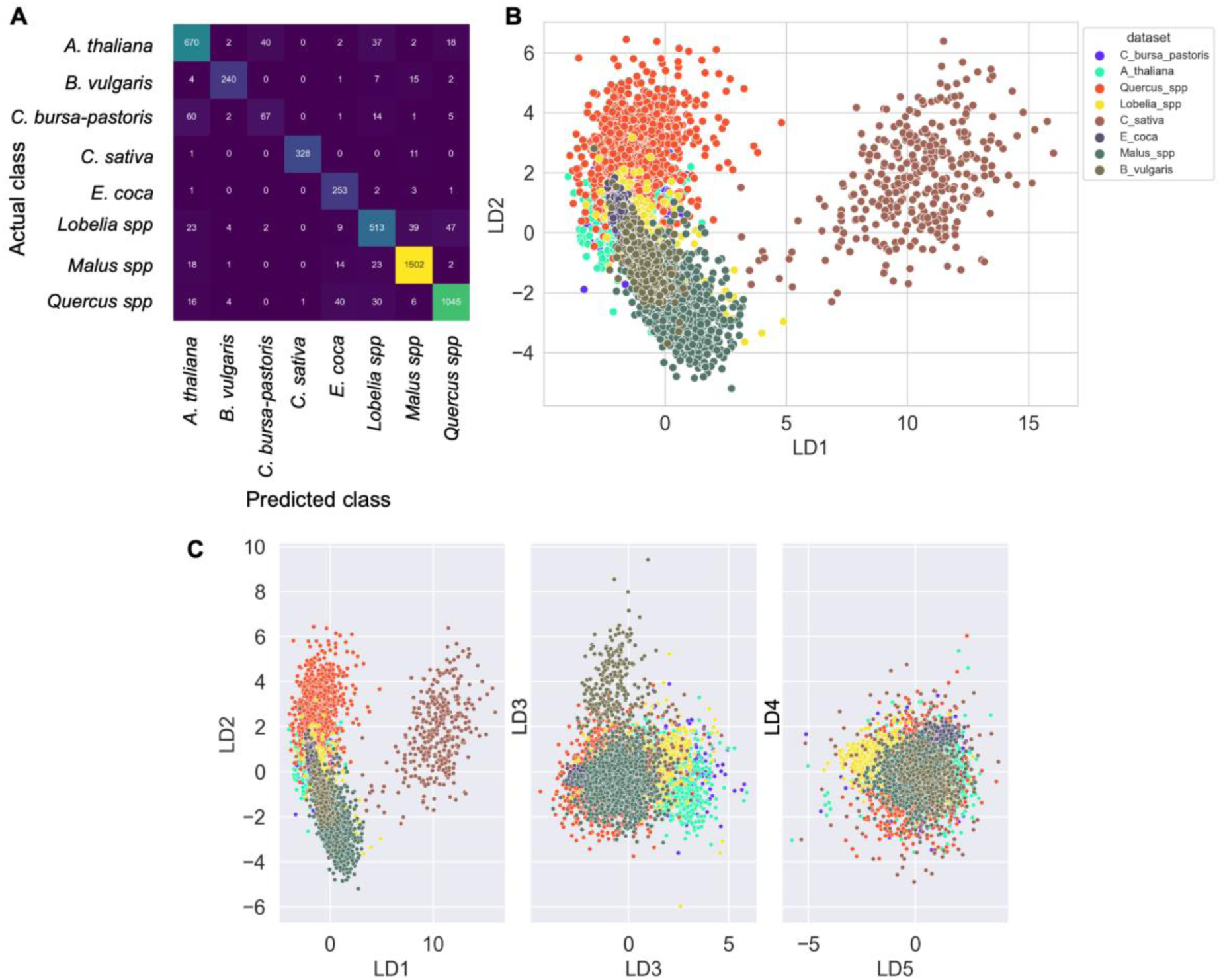
Linear discriminant analysis of all leaves. **A**. A confusion matrix of actual and predicted leaves from each species dataset based on the 100-pseudo landmarks. **B**. A scatterplot of predicted leaves from each species dataset colored by each species dataset. **C**. scatterplots of predicted leaves from each species datasets from LD1 to LD5, colored by each dataset.

### Modeling leaves across development and by genetic data reveal greater insights into the mechanisms for shape change

In addition to dimensional reduction techniques like PCA and LDA, leaves from plants that grow in a continuous series can be modeled using their growth positions along the stem. Modeling leaves in a continuous series provides insights into how leaf shape changes across development, which is directly related to changes in gene expression. An example of this continuous modeling can be seen in leaves from *Passiflora* or passion fruit/maracuyá species. To demonstrate a change in leaf shape from the base of the shoot (the earliest emerged leaves) to tip (the latest emerged leaves), we used previously published data (Chitwood and Otoni, 2017a, b) to model leaves by the relative node position. 3,299 leaves in total are analyzed. The total number of leaves/nodes per plant often varies, which may place leaves of the same node number in different developmental stages. Therefore, to compare growth order and leaf development, it is necessary to contextualize the leaf growth order for each plant using relative node number, or the node number counting from the shoot base divided by the total number of nodes.

We modeled leaf shape across development as each x and y coordinate value for each pseudo-landmark as a function of relative node number for each shoot using second degree polynomials. Using these continuous models, we then visualized 40 leaves in order from the shoot base to tip for each *Passiflora* species model, colored by previously-defined *Passiflora* leaf morphotypes (Fig. 6A). Morphotypes A, E, F, and G show little variation in leaf shape across development. In contrast, morphotypes B, C, and D show more variation across development, where the leaves closer to the base are more uniform than the leaves closer to the tip of the shoot. When we compare individual *Passiflora* leaves using a PCA morphospace, we see significant separation between morphotypes (Fig. 6B), with overlap of morphotypes E, F, and G, representing leaves that are more entire. Morphotypes A and B overlap in PC space with eigenleaf representations that are wider than they are long, with no obvious leaf tip point (Fig. 6B). Morphotype C separates on its own and is represented by a tri-lobed leaf morph. Morphotype D does not show distinct separation from any other morphotype and can be found in all three clusters. When we compare developmental models of *Passiflora* leaves in PC space using continuous curves that represent modeled leaves as a function of relative node, we see a similar separation of morphotypes into three distinct groups (Fig. 6C). In general, the first leaves for each morphotype and species are less lobed and more entire, originating from the center of the morphospace (Fig. 6C). Later leaves then include more shape diversity, with variations in overall leaf margin architecture (Fig. 6C). By modeling changes in shape across development, we can understand when and where during development leaf shape varies across species.

**Figure six.**
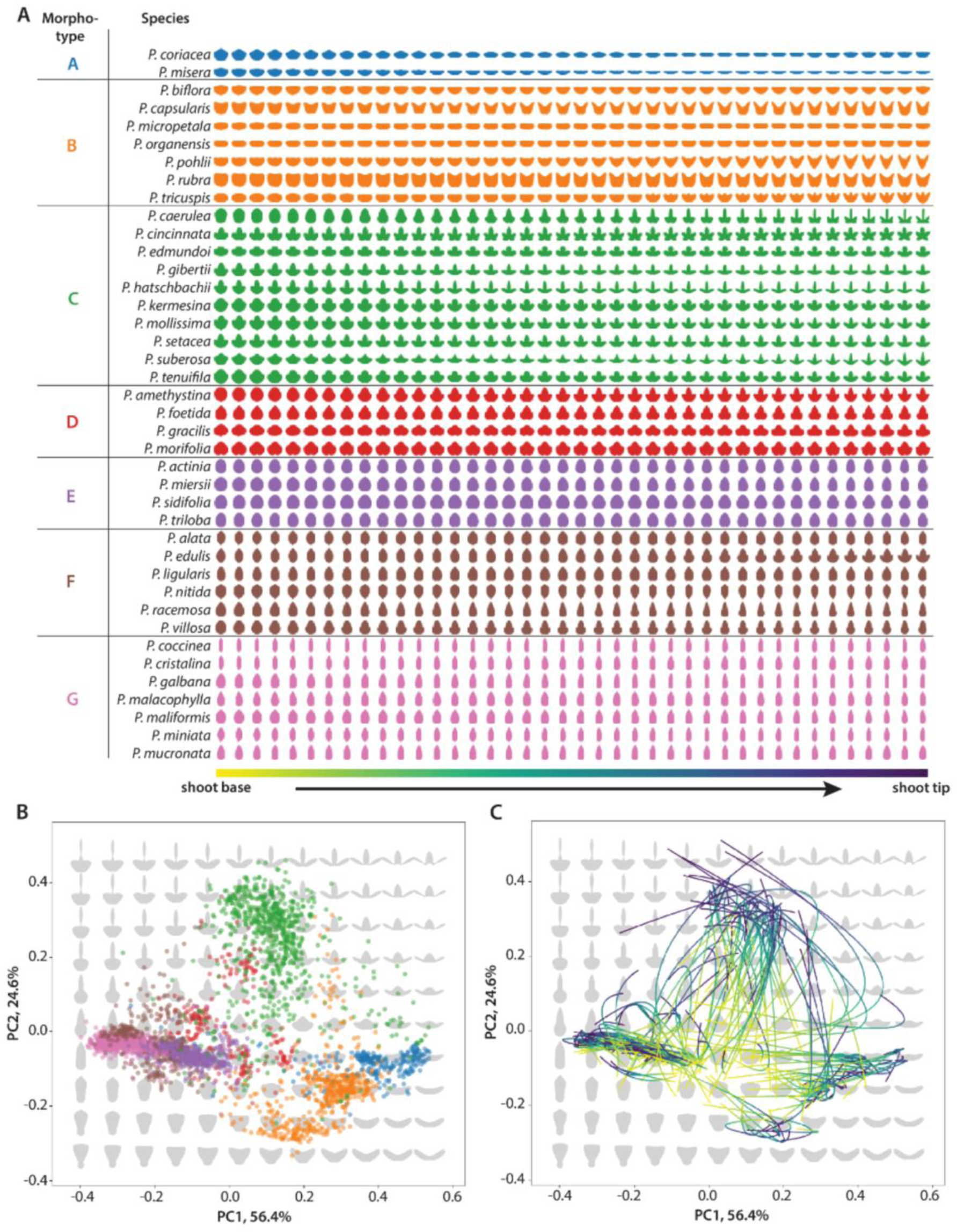
Continuous models of leaf development across nodes. **A**. For each of 40 *Passiflora* species, organized and colored by morphotype, the average developmental model across 40 intervals, organized from shoot base to shoot tip. **B**. A morphospace for individual leaves colored by morphotype. **C**. Developmental models for each plant projected back onto the morphospace. Each curve represents a developmental model for a single plant, colored using a viridis color scale from the shoot base to shoot tip.

In addition to the other leaf species included in both our collaborative dataset and the previously published *Passiflora* data, the compound tomato (*Solanum lycopersicum*) leaf has been used to study the genetic and developmental basis of leaf shape. Previously published leaf shape data (Chitwood et al., 2013) from 76 *S*. *lycopersicum* introgression lines (ILs) that feature small introgression from the wild tomato relative *S. pennellii* into a *S*. *lycopersicum* background, reveal genetic differences in shape between the terminal, proximal left, and distal right lateral leaflets. 11,276 leaflets were analyzed. Because each IL contains a defined *S. pennellii* introgression, a significant statistical difference in a trait between an IL and the *S. lycopersicum* background indicates its genetic basis (the introgression).

Chitwood, et al,. 2013 showed that it is possible to map QTL’s (quantitative trait loci) to overall shape descriptors or principal component summaries of leaflet shape. Here, we continuously map each point across the leaflet individually. By continuously mapping points across the leaflet, we can identify which QTL’s are associated with continuous shape change across leaflets. To convert pseudo-landmarks into a continuous trait value, we take the Euclidean distance of a pseudo-landmark of a sample to its counterpart on the overall mean, where values outside the mean are positive and inside are negative in value. We use a Spearman’s rank correlation to correlate the presence or absence of an introgression bin with the displacement of each pseudo-landmark from the overall mean of the terminal or right or left distal lateral leaflet. We can visualize the degree of displacement of each pseudo-landmark from its respective mean leaflet or as a heatmap, plotting pseudo-landmarks along the leaflet (leaf region) as a function of the genetic map (chromosomal introgressions) and coloring by the statistical strength of the interaction and its direction (Fig. 7 Top – Bottom). Results of our continuous mapping reveal the regions of the terminal, lateral right, and lateral left leaves that correspond with previously described QTLs (Chitwood et al., 2013). Most importantly, we can now identify which regions of which leaflets the previously described QTLs (represented by introgression lines, or ILs) are acting on. IL2-1 exhibits narrower leaves proximally at the base of the lateral leaflets, transgressive beyond *S*. *lycopersicum.* IL9-1-2 exhibited narrower leaves distally at the tip of terminal and lateral leaflets, again transgressively beyond that of *S*. *lycopersicum.* IL4-3, one of the most extreme tomato ILs, exhibited wider leaves especially in distal regions of the lateral leaflets, but also somewhat in the terminal leaflet, similar to *S. pennellii*. IL5-4, exhibited wider leaves specifically in the terminal leaflet, and proximally at the base. Therefore, through the use of simple correlation, pseudo-landmarks, and a defined genetic resource we are able to model the underlying genetic architecture associated with specific parts of tomato compound leaves.

**Figure seven.**
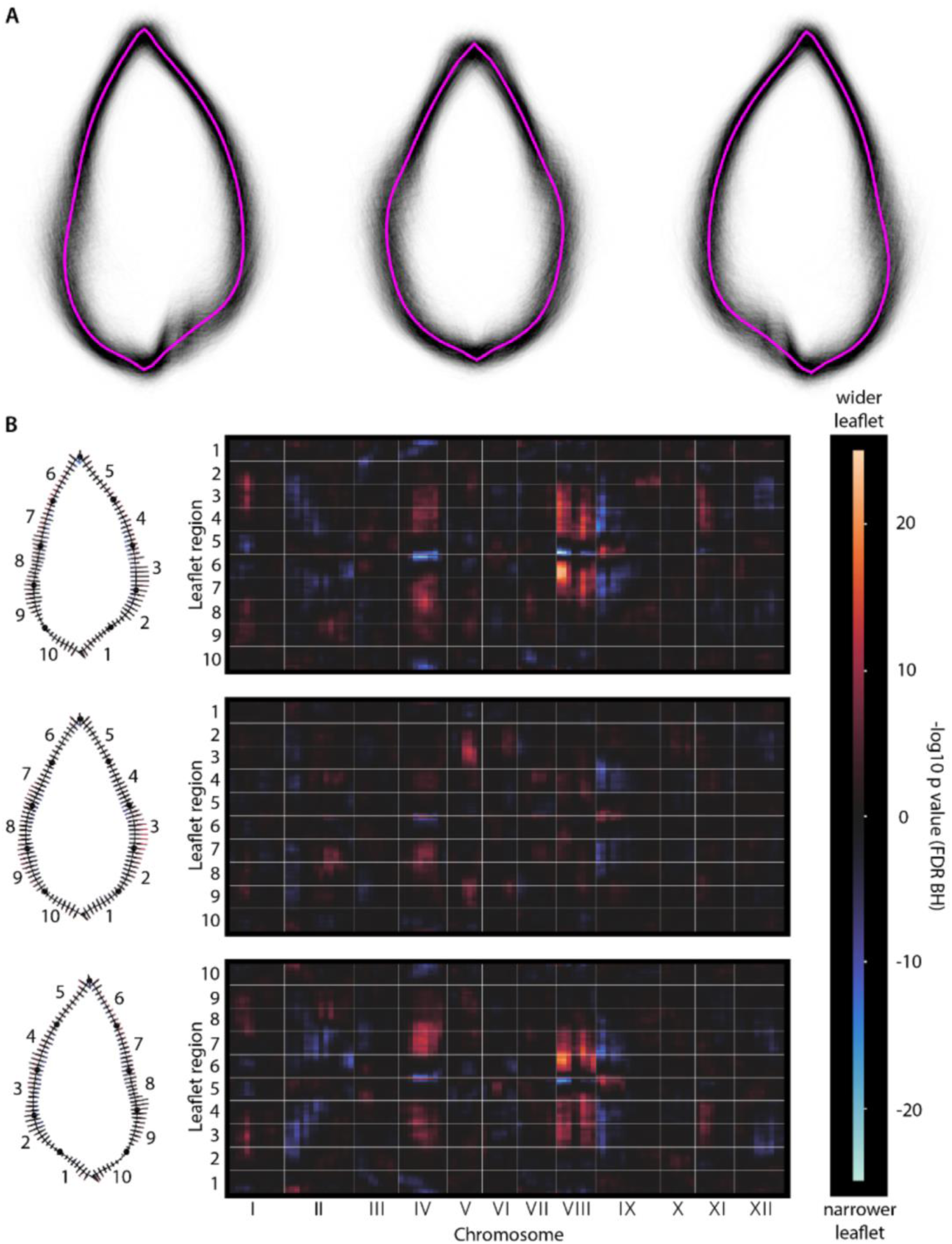
The mapping of tomato pseudo-landmark positions relative to mean leaflet shape to genetic loci. **A**. Left to right: left distal lateral, terminal, and right distal lateral tomato leaflets. The generalized Procrustes analysis (GPA) mean for each is shown in magenta and superimposed leaflet outlines are transparently depicted in black. **B**. The genetic basis of leaflet pseudo-landmarks. Top to bottom: left distal lateral, terminal, and right distal lateral tomato leaflets. Left, pseudo-landmarks are assigned to ten regions along the leaflet outline, indicated by number. For quantitative trait loci with FDR BH-adjusted p values less than 0.05, a line segment normal to the associated mean pseudo-landmark position and colored by p value and direction is shown, showing the genetic effects on shape in specific leaflet locations. Right, for each leaflet type a heat map, colored by -log base 10 FDR BH-adjusted p values, for each leaflet region (Arabic numerals) by genetic bins across chromosomes (Roman numerals).

## Discussion

### Advantages of pseudo-landmark and Procrustes based analysis

As demonstrated above, one of the key advantages of using pseudo-landmarks and Procrustes based leaf shape analysis is the ability to include highly variable and irregular leaf shapes. Geometric morphometric analyses that include landmarks rely on biological homology between species and shapes. Additionally, traditional landmarks are typically limited in number and are often hand selected. However, for many species, the level of biological homology necessary for true landmarks makes traditional geometric morphometric analyses incredibly difficult. For example, *C. bursa-pastoris* leaves exhibit incredible variation in lobe number, lobing angle, and lobe depth making finding homology between leaves nearly impossible. In contrast, by leveraging the two landmarks common to all leaves—base and tip points—and by using equidistantly placed pseudo-landmarks between these anchors, we can not only increase the number of comparable datapoints, but also align datapoints to infer new points of homology between species and shapes. The ability to align and compare shapes cannot be overstated.

When modeling changes in shape, determining a mean shape and the displacement of individuals from the mean is key for annotating and measuring changes in shape. Both the Procrustes distance and the combinations of pseudo-landmarks can be used to determine the differences between shapes using linear modeling. Additionally, combinations of pseudo-landmarks show the locations along the leaf margin that are most responsible for variation between leaves, revealing potential tissue-based locations for selection and phenotypic plasticity, and creating features that capture additional leaf shape complexity that is lost if only dimensions or other summary features are considered. Overall, using a pseudo-landmark and Procrustes based approach is an easy, fast, and informative analysis for measuring the variation in leaf shape.

### Leveraging dimension reduction techniques and modeling

Another advantage of using pseudo-landmarks to describe and measure leaf shape is that pseudo-landmarks can be used as an input for both PCA and LDA, to create visualizations of morphospaces and predictive models, respectively. By creating a PCA morphospace, not only can we recreate theoretical leaf shapes that continuously define variation between leaves, we can measure and visualize the relationships between leaves in our dataset across multiple, orthogonal axes. As previously described, there are boundaries to the PCA morphospace (Chitwood and Mullins, 2022) that inform the possible constraints in leaf shapes with respect to development. This can be observed towards the fringes of the morphospace far from the center of the dataset, in which eigenleaves stretch or fold back upon themselves in ways that actual leaves never do. This phenomenon is also seen for leaf shapes that pseudo-landmark methods have difficulty in capturing, for example very thin leaves or leaves with numerous and irregular lobes. Such difficult shapes nonetheless represent extreme leaf morphs that in reality are far from the common, ancestral leaf ideotype. Similarly for LDA, by using linear combinations of pseudo-landmarks, those parts of the leaf that most contribute to the separation of shapes can be used for prediction. The classification resulting from an LDA model likewise is informative of the relationship of individual leaf shapes to each other, with leaf shapes more similar to each other and the overall mean leaf more likely to be confused with each other, whereas more unique and extreme leaf shapes will be easily predicted.

### Biological relevance of landmarks and shape

Just as pseudo-landmarks precisely convert specific positions along the leaf margin into features that can be compared across all leaves to determine those that most vary by genetic or developmental factors, so too can they be used to identify aspects of leaf morphology with respect to function. Paleobotanical and common garden studies have thoroughly demonstrated that leaf shape variation affects numerous processes necessary for plant fitness in various environments. Physiological studies show that deeply lobed leaves have lower ratios of mesophyll tissue to large conductive veins which lowers the resistance of water transport throughout the leaf (Xu et al., 2009; Ding et al., 2020; Sakurai and Miklavcic, 2021). In less lobed leaves, smaller veins increase water resistance because water must travel laterally between cells (Ding et al., 2020; Sakurai and Miklavcic, 2021). Additionally, heat convention is more rapid in small leaves than in large leaves and the thickness of the boundary layer increases with leaf length, creating greater disparities in heat dissipation (Bednarz and van Iersel, 2001; Peppe et al., 2011). Deeply lobed leaves decrease the distance across the lamina that heat must travel, resulting in greater heat transfer in lobed leaves than in entire leaves (Zwieniecki et al., 2004). As such, the biological trade-off between leaf structure and function has been well documented (de Boer et al., 2016; Gutschick, 2016). Therefore, by knowing which parts of leaves vary between scales, we can infer the evolutionary and physiological function that may result from leaf shape variation.

### Future uses for pseudo-landmark and Procrustes based analysis

As pseudo-landmark and Procrustes based analyses become more widely applied to leaf shape, future uses for this analysis include creating synthetic leaves (Balant et al., 2024; Chitwood et al., 2025), or theoretical eigen leaves that sample representations across the morphospace along increasing numbers of PCs, that can be used to infer unmeasured continuous variation in leaf shape. As more leaves are measured through automated segmentation of leaf shapes from imaged leaves from common gardens and herbaria, we will be able to predict shape boundaries and visualize theoretical leaf shapes across biologically relevant axes. By creating developmental models, we will similarly capture continuous developmental variation, which from previous studies appears to be additive and orthogonal to differences in leaf shape between species. For example, by modeling leaves across nodes, whether shoots or rosettes, we can generate continuous models of leaf shape as a function of plastochron index, which measures the time interval between the formation of two successive leaves. The plastochron index describes the age of the plant based on morphological traits like leaf shape, and additionally can be used to predict both developmental trajectories of heteroblasty and ontogeny across nodes. The morphometric featurization of leaf shape by pseudo-landmarks can also be used for high-resolution genetic studies. Pseudo-landmarks themselves and dimension reduction outputs like PCs can be used directly in genome wide association studies (GWAS) or analyses with RNASeq. However, by measuring the Euclidean distance displacement of pseudo-landmarks from their respective means as we have shown here, the genetic basis of individual points can be independently tested from each other. Further, the Procrustes distance itself, a measure of overall similarity between two shapes, can be used to measure within and between group variation relative to respective GPA means. Finally, the pseudo-landmarks and any combination of the other measures mentioned here can be used in deep learning, where pseudo-landmarks can be used to create artificial neural networks that work to understand leaf shape pattern recognition at a deeper level. The estimation of a morphospace defined by distances between samples, combined with similar spaces defining spaces of molecular profiles, opens up the possibility of using graph learning to predict phenotype from genomic information, leveraging the projected position of samples onto previously defined landscapes.

### A call to action: building a morphospace of all leaf shapes

In this study, we have shown that examining leaf shape variation can be easy, cheap, and fast. Importantly, by using base and tip points that are common and homologous to all leaves and placing equidistantly spaced pseudo-landmarks on each side of the leaf, any leaf can be compared in overall similarity to any other. Therefore, as we continue to study leaf shape as a wider plant science community, we would like to challenge any researchers that are interested in leaf shape to not only use the analyses and data presented in their study with these analyses techniques, but to combine your data with the leaf shapes shared here to create a continuously growing morphospace of all leaves. Already segmentation algorithms applied to digitized herbarium samples promise millions of leaves from across species and environments to be available to compare leaf shapes from more narrow studies to be compared against all leaves. Studies that sample genetic, developmental, physiological, or evolutionary processes in depth will be able to combine this valuable information against the landscape of all leaf shapes. An unprecedented time to realize a genetic, developmental, evolutionarily, and functional synthesis of leaf morphology across plants is upon us.

## Supporting information

Table one

Table two

## Data availability

The leaf outlines and individual dataset metadata can be found in the GitHub repository (DOI: 10.5281/zenodo.15685723) for this work. The Jupyter notebook and accompanying code use in this leaf shape analysis can also be found in the GitHub repository for this work.

## Contributions

AH collected the experimental data from each dataset group, collected experimental data using *C. bursa-pastoris*, created the analysis, prepared the leaf shape analysis including the Jupyter notebook and GITHUB repository, prepared and edited the manuscript, and submitted the manuscript for review. DHC prepared the initial leaf shape analysis and consulted on further analysis, reviewed and edited the manuscript, and contributed to submitting the manuscript for review. AC, CB, SH, and SW generated leaves from the *Lobelia spp.* dataset and reviewed and provided feedback on the manuscript. CH and CH generated leaves from the *Quercus spp*. dataset and provided feedback on the manuscript. JH, LS, ZM generated leaves from the *Malus spp*. dataset and provided feedback on the manuscript. AH and RUC generated leaves from the *A. thaliana* dataset and provided feedback on the manuscript. MB and AP generated leaves from the *C. sativa* dataset and provided feedback on the manuscript. RN and AD generated leaves from the *B. vulgaris* dataset and provided feedback on the manuscript. EJ, CH, CC, EA, GD generated leaves from the *C. bursa-pastoris* dataset and provided feedback on the manuscript. YASR and ER generated leaves from the *E. coca* dataset and provided feedback on the manuscript.

